# Improved imputation of summary statistics for admixed populations

**DOI:** 10.1101/203927

**Authors:** Sina Rüeger, Aaron McDaid, Zoltán Kutalik

## Abstract

**Motivation:** *Summary statistics imputation* can be used to infer association summary statistics of an already conducted, genotype-based meta-analysis to higher genomic resolution. This is typically needed when genotype imputation is not feasible for some cohorts. Oftentimes, cohorts of such a meta-analysis are variable in terms of (country of) origin or ancestry. This violates the assumption of current methods that an external LD matrix and the covariance of the Z-statistics are identical.

**Results:** To address this issue, we present variance matching, an extention to the existing *summary statistics imputation* method, which manipulates the LD matrix needed for *summary statistics imputation*. Based on simulations using real data we find that accounting for ancestry admixture yields noticeable improvement only when the total reference panel size is > 1000. We show that for population specific variants this effect is more pronounced with increasing *F*_*ST*_.

## 1 Introduction

Genotype data for genome-wide association studies (GWASs) are often collected using DNA chips, which cover only a small fraction of the variable genome. To be able to combine GWASs that measured different sets of genetic markers (due to differences in the content of commercial arrays), genetic information has to be inferred for a common set of markers. Such inference exploits the fact the neighbouring SNVs are often in linkage disequilibrium (LD), which has been well-quantified in different human populations. Statistical inference of these untyped SNVs in a study cohort, therefore, relies on an external reference panel of densely genotyped or sequenced individuals. The inference process is termed *imputation*, of which there are two main types. *Genotype imputation* (Marchini and Howie, 2010) first estimates all haplotypes both in the reference panel and the study cohort, then using a Hidden Markov Model every observed haplotype in the study cohort is assembled as a probabilistic mosaic of reference panel haplotypes. The reconstruction facilitates the computation of the probability of each genotype for every SNV of the reference panel in each individual of the study cohort. Having imputed the genotype data set, one can then run an association scan with an arbitrary trait and obtain association summary statistics. *Summary statistics imputation* Pasaniuc et al. (2014) on the other hand starts off with association summary statistics available for all genotyped markers and infers, combined with a reference panel, directly the association summary statistics of SNVs available in the reference panel. More specifically estimating the local pair-wise linkage disequilibrium (LD) structure of each genetic region using the reference panel and combining it with association summary statistics allows to calculate a conditional expectation of normally distributed summary statistics. This latter approach is the central focus of our paper. Compared to *genotype imputation*, *summary statistics imputation*is much less demanding on computational resources, and requires no access to individual level genetic data.

Methods making use of summary statistics, such as calculating genetic correlation (Bulik-Sullivan *et al*., 2015), approximate conditional analysis (Yang *et al*., 2012) or causal inference (Burgess *et al*., 2013), have gained interest in recent years, because they bypass the need of genotype data, but mimic it by making use of external reference panels. These methods could profit from summary statistics being available on an arbitrarily chosen panel of SNVs - provided by *summary statistics imputation*. However, it is not clear how to optimally combine different LD reference panels for summary statistics emerging from a meta-analysis of a large number of different studies (coming from different countries/regions), with potentially different ancestries. To ensure accurate imputation of such “admixed” meta-analyses, we propose a method called *variance matching* that, for each genomic region, optimally combines reference panels to best match the local LD pattern of the underlying GWAS population. Using a simulation framework, we compare *variance matching* to a *benchmark* solution and one of previously proposed approaches (Lee *et al*., 2015a,b; Park *et al*., 2015).

## 2 Methods

### 2.1 Summary statistics imputation (SSimp)

We assume a set of univariate effect size estimates ***a***_*i*_ are available for SNVs *i* = 1,…, *I* from a linear regression between a continuous phenotype ***y*** and the corresponding genotype ***g***^*i*^ measured in *N* individuals. Without loss of generality we assume that both vectors are normalised to have zero mean and unit variance. Thus 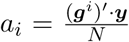 and 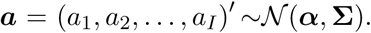 Σ represents the pairwise covariance matrix of effect sizes of all *i* = 1,…, *I* SNVs.

To estimate the univariate effect size *α*_*u*_ of an untyped SNV *u* in the same sample, one can use the conditional expectation of a multivariate normal distribution. The conditional mean of the effect of SNV *u* can be expressed using the effect size estimates of the tag SNVs (Eaton, 1983; Pasaniuc *et al*., 2014):

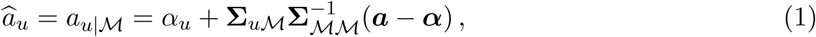

where 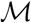 is a vector of marker SNVs, 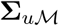 represents the covariance between SNV *u* and all 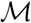 markers and 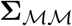 represents the covariance between all 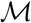 markers.

We assume that estimates for the two covariances are available from an external reference panel with *n* individuals and denote them 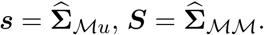 The corresponding correlation matrices are *c* =*N* · *s* and ***C*** = *N* · ***S*** (with *γ* and 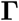 as the corresponding true correlation matrices). Further, by assuming that SNV *u* and the trait are independent conditioned on the 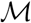 markers, i.e. 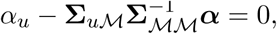, Eq. (1) becomes

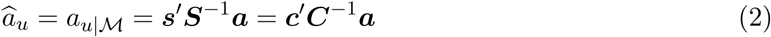

One can also choose to impute the Z-statistic instead, as derived by Pasaniuc *et al*. (2014):

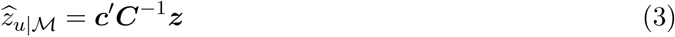

with 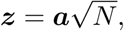 when the effect size is small (as is the case in typical GWAS).

Similar to Pasaniuc *et al*. (2014), we chose 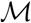 to include all measured variants within at least 250 Kb of SNV *u*. To speed up the computation when imputing SNVs genome-wide, we apply a windowing strategy, where SNVs within a 1 Mb window are imputed simultaneously using the same set of 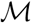 tag SNVs the 1 Mb window plus 250 Kb flanking regions on each side.

Summary statistic imputation in the context of genomic data was first described by Wen and Stephens (2010), where they inferred allele frequencies for an untyped SNV by a linear combination of observed allele frequencies (BLIMP). Lee *et al*. (2013) then further extended the method to the application of linear regression estimates (DIST). Later, Pasaniuc *et al*. (2014) included a sliding window, which allowed partitioning of the genome into smaller pieces to facilitate imputation on a larger scale, and introduced a different shrinkage approach (IMP-G). Since then a few extensions have been published, that mainly concentrate on summary statistic imputation for admixed populations (Lee *et al*., 2015a,b; Park *et al*., 2015) or including covariates (Xu *et al*., 2015).

#### Shrinkage of SNV correlation matrix

To estimate *C* (and c) we use an external reference panel of *n* individuals. Since the size of *C* often exceeds the number of individuals (*q* ≫ *n*), shrinkage of matrix *C* is needed to guarantee that it is invertible. By applying shrinking, the modified matrix *C* becomes

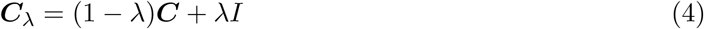

Even though *c* is not inverted, we still shrink it to curb random fluctuations in the LD estimation in case of no LD.

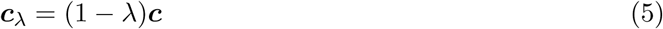

Inserting *c*_λ_ and *C*_λ_, Eq. (2) then becomes

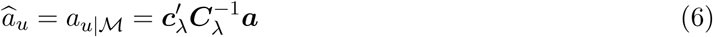

Note that λcan vary between 0 and 1, with λ = 1 turning *C* to the identity matrix, while λ = 0 leaves *C* unchanged. Here, we mainly focus on λ changing with the reference panel size 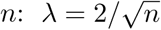 (Lee *et al*., 2014a).

### 2.2 Optimal combination of reference panel subpopulations to match the GWAS sample

For *summary statistics imputation* we would like to estimate the local LD structure of each region in the GWAS population 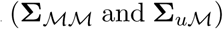 and to do so we use a (sequenced) reference population, yielding estimates *C* and c. Clearly, the closer these estimates are to the real values, the better the imputation will be (i.e. smaller the estimation error in Eq. (6)). Our aim is to find a weighted mixture of the reference sub-populations that has an LD structure as similar as possible to the LD in the GWAS population.

Park *et al*. (2015) proposed an elegant, generalised approach to weight population LD structure. Their algorithm *Adapt-Mix* choses weights (*w*^*am*^) based on optimising an objective function. In the case of imputation the objective function is the MSE of the (re-)imputed Z-statistics at observed SNVs. Lee *et al*. (2015a) developed *Distmix*, which minimises the Euclidean distance between allele frequencies of the reference panels and the GWAS study, but ignores the variance-bias trade-off.

While the true LD structure of the actual GWAS population is rarely known, the GWAS allele frequencies are routinely calculated (even if not always reported for out-dated privacy preserving reasons) in meta-analytic studies. In the following we show how this information can be exploited to improve *summary statistics imputation*.

First, suppose that the reference panel is made up of *P* subpopulations of sizes *n*^(1)^, *n*^(2)^, …, *n*^(P)^. Next, we introduce a set of weights ***w*** = (*n*^(1)^, *n*^(2)^, …, *n*^(*p*)^)/*n* = *w*_1_,*w*_2_, …, *w*_*p*_, which can be viewed as the collection of weights that determine the reference population mixture, i.e. 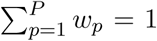 and *w*_*p*_ ≥ 0.

We can calculate the covariance *s* as a function of these weights (*s*(*w*)),i.e. for each subpopulation we calculate the covariance separately (*s*(*p*)), and then combine them, weighted by their weights ***w***:

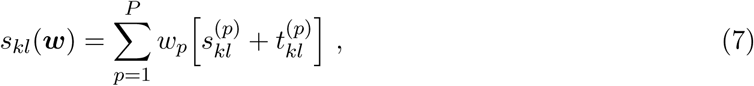

where 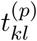 is the between-group covariance for variants *k* and *l* in population *p*:

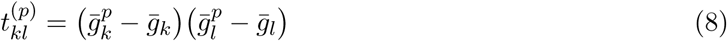

and 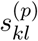 denotes the covariance for variants *k* and *l* in population *p*:

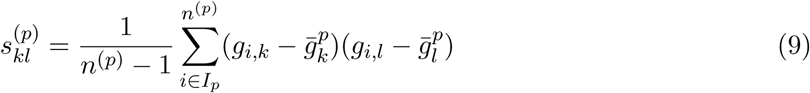

*g*_*i,k*_ refers to genotype of variant *k* for individual *i*. The overall, mean population genotype dosage (i.e. twice the allele frequency) is naturally defined as the weighted mean sub-population genotype dosage 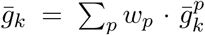 and 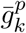 being the average genotype dosage in population 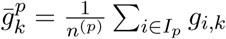 and *I*_*p*_ refers to the indices of individuals contained in population *p*.

While the reference panel population sizes are being fixed at *n*^(*p*)^ and we defined *w*_*p*_ ∝ *n*_*p*_, we could use any arbitrary weights ***w*** in order to match a GWAS population, which has different population proportions than the reference panel. This manipulation of the covariance estimation can be used to adapt the reference panel population structure towards the population structure that is observed in GWAS summary statistics.

The corresponding correlation between SNV *k* and *l* from a reference panel with specific chosen weights ***w*** is

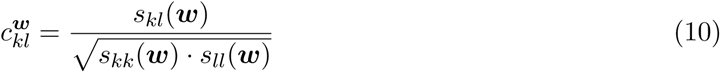

Our goal is to quantify the mean squared error (MSE) between the true GWAS LD matrix 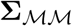 and the LD matrix estimated from the reference panel 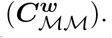 Since we cannot estimate the off-diagonal values of the GWAS covariance matrix, we focus on its diagonal elements and estimate them from the GWAS allele frequencies. The MSE of Eq. (7) for SNV *k* can be written as

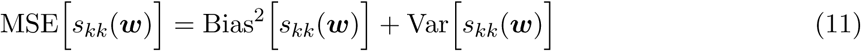

In short, the MSE of Eq. (7) depends on known quantities (mean genotype dosage 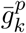 for SNV *k* in population *p*; sample sizes *n*^(*p*)^ of the reference panel population *p*; average genotype for SNV *k* in the GWAS study: 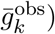 and the unknown mixing parameter ***w***. Assuming Hardy-Weinberg equilibrium (HWE), we showed that the variance term is a sixth-degree, while the squared bias is a fourth-degree polynomial in ***w***. Details to derivations of the MSE are provided in Supplement A.2.

We aim to find a ***w*** that minimises the MSE in Eq. (11) for a set of M SNVs, for which we know the allele frequencies 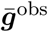 in the GWAS population and can estimate 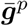 from a reference panel:

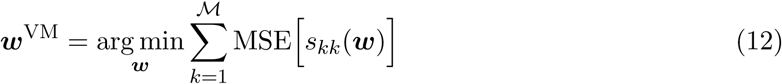

with 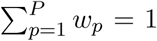 and *w*_*p*_ ≥ 0. We call this method *variance matching* (vm), as we are using the GWAS allele frequencies to match genotype variances. Parameter ***w***^VM^ gives us an estimation of the population weights used in Eq. (7).

Finally, we substitute ***w***^VM^ into Equation Eq. (10) and plug it into Eq. (6):

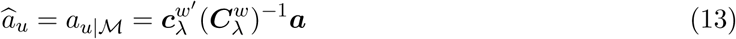

Detailed derivations of Eqs. (7) to (12) can be found in Supplement A.

### 2.3 Reference panels

As reference panels we used genetic data from the 1000 Genomes Project Consortium (2010).

### 2.4 Simulation

#### 2.4.1 Simulation of GWAS summary statistics

For simulation of GWAS summary statistics we used data from the five European subpopulations `CEU, GBR, FIN, TSI` and `IBR` of the 1000 Genomes project (1KG). We chose to up-sample chromosome 15 using HAPGEN2 (Su *et al*., 2011) to 5′000 individuals for each subpopulation, yielding a total of 25′000 individuals. Of these 5000 individuals per population we used half each to generate a GWAS with an *in silico* phenotype. The remaining 12′500 individuals were used as reference panel for summary statistic imputation.

We split chromosome 15 into 74 disjoint regions of 1.5 Mb. Due to the sliding window imputation approach we did not include regions at the very start and end of the chromosome. In each region we chose a causal variant ***g*** randomly from all SNVs with minor allele frequency between 0.05 and 0.2. We simulated an *in silico* phenotype ***y*** using a normal linear model ***y*** = *β****g*** + *ε* with 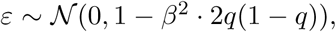 where *q* is the allele frequency of the causal SNV and *β* was selected such that the explained variance *β*^2^ · 2*q*(1 - *q*) is set to 0.02. To obtain the association summary statistics we ran linear regression for each variant *k* in the 1.5 Mb region, yielding effect size and standard error estimates *a_k_, se(a_k_)*, from which we calculated the standardised effect size estimate 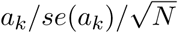 (*N* being the sample size).

#### 2.4.2 Applying summary statistics imputation and comparing methods

We constructed GWAS genotype datasets with a fraction *w*^+^ of Finnish individuals. The total number of individuals in the GWAS genotype dataset was constant at 2′500. Next, we calculated a re-weighted *C* from our reference panel (with weight ***w***). We then created different scenarios by repeating this procedure for many different GWAS compositions (i.e. we varied the Finnish fraction *w*^+^ between 0 and 1 in 0.2 increments) and weights ***w*** of Finnish for the correlation matrix of the reference panel (which we varied between 0 and 1 in 0.05 increments). For each scenario, we calculated three MSE for a set of imputed SNVs (Eq. (14)): first, the MSE of the standardised effect size; second, for the *variance matching* approach we calculated the MSE of (the diagonal of) matrix *C* estimated from the reference panel; third, for the *Adapt-Mix* approach we calculated the MSE of the standardised effect sizes of observed SNVs, as described in Park *et al*. (2015).

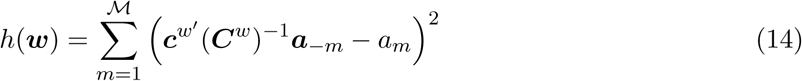

By minimising each error measurement over all ***w***, the first MSE will determine ***w***\*, which gives the theoretically best possible solution.

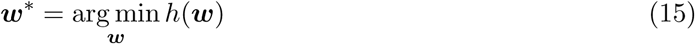

In our approach, the estimated MSE of matrix *C* will determine ***w***^VM^. While in the best competing algorithm, Adapt-Mix, the minimised MSE of the reimputed Z-statistics determines the value of ***w***^AM^. We chose to vary the proportion of the Finnish population, as it differs the most from other populations in Europe in terms of allele frequencies and LD structure Lim *et al*. (2014), McEvoy *et al*. (2009). The remaining four populations of the European 1000 Genomes populations share equal weights in all scenarios. In our simulation we are looking at HapMap SNVs only as tag SNVs. There are between 167 and 1103 tag SNVs per region with mean 635. We imputed on average 1′743 SNVs per region (out of 74 in total).

## 3 Results

*Summary statistics imputation* works through combining summary statistics from a set of SNVs with pairwise SNV LD information obtained from an external reference panel. We extended the most recent summary statistics method (Pasaniuc *et al*., 2014) by an optimal assembly of the LD matrix from a mixture of reference panels. Because our approach optimises the diagonal of the covariance matrix, we term our method *variance matching*. We compare our it to *Adapt-Mix* by Park *et al*. (2015) and a *benchmark* solution.

### 3.1 Simulation framework

To assess *variance matching* we used upsampled datasets, yielding 25′000 European individuals in total. GWASs were simulated using *in silico* phenotypes. This semi-simulation framework allowed us to study the impact of the reference panel sizes (up to 12′500) and their composition.

In brief, for various ancestry compositions of simulated GWAS sample we computed association summary statistics, masked a fraction of SNVs and imputed them. When imputing a single SNV we used tag SNVs within at least 250 Mb. For the imputation of an entire region we used a sliding window of 1 Mb with 250 Kb flanking regions on each side.

More specifically, for each simulated GWAS, we fixed the proportion of the Finnish subpopulation of the European reference panel of 1000 Genomes Project Consortium (2010) in the GWAS, then let the proportion of this population vary in the reference panel used for LD estimation. We repeated this for different Finnish proportions in the GWAS and in the reference panel (varying from 0 to 1), calculated each time the MSE between the estimated and the imputed standardised effect sizes (*h(w)*, Eq. (14)) and determined the *benchmark* weight that yields a minimal MSE (denoted as ***w***\*,Eq. (15)). In parallel, we applied for each fixed proportion of Finnish in the GWAS the *variance matching* and the *Adapt-Mix* approach to determine their optimal weight — ***w***^VM^ and ***w***^AM^ — in the reference panel (Figure S1). To identify other factors that influence the choice of weights, we grouped the 637′153 SNVs into population specific (76′013) and population non-specific (561′140) groups (based on *F*_*ST*_ ≥ 1% vs *F*_*ST*_ < 1%, respectively), and ran the simulation from small to large reference panels (*n* = {500, 1′000, 2′500, 5′000,12′500}).

### 3.2 Improving summary statistics imputation via variance matching

Ultimately, we are interested in two comparisons. First: the optimisation of weights versus the ad-hoc reference panel (which has roughly equal weights in the European sub-panel, i.e. ***w*** = 0.2). Second: how *Adapt-Mix* and our notvel method *variance matching* perform compared to the *benchmark* estimation (the best possible choice if we were to know the true effect size). These two comparisons are presented in Fig. 1, where we compare the MSE of the three optimal weights (***w***^VM^, ***w***^AM^, ***w***\*) determined by each method relative to the MSE when using equal weights: MSE-ratio = *h*(***w***)/*h*(***w*** = 0.2), with *h* denoting the MSE between the estimated and the imputed effect sizes described in Eq. (14).

**Figure 1.**
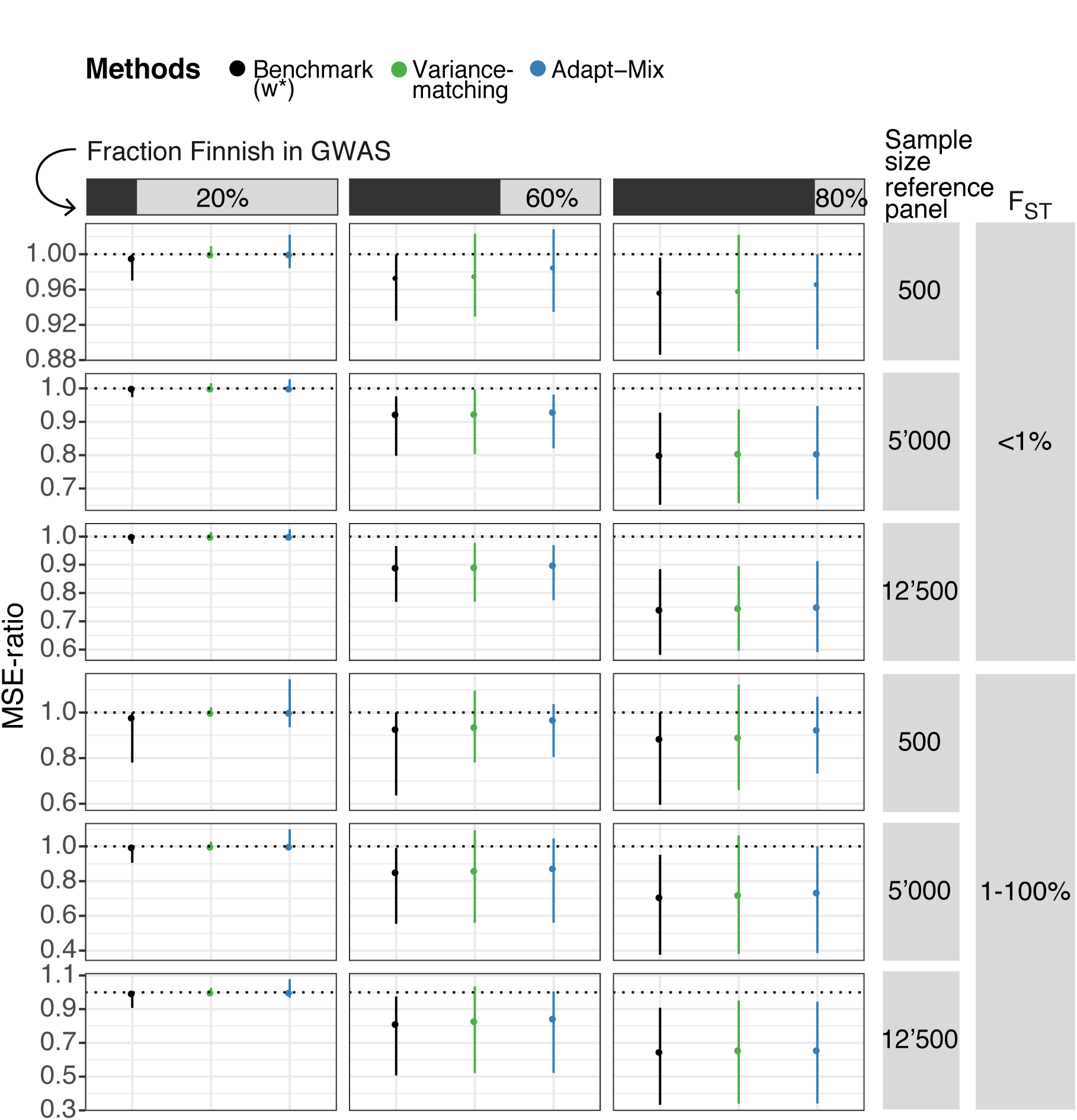
Comparison of methods accounting for population structure. This figure shows the comparison of each *w*-optimisation method with respect to choosing the full European panel of 1000 Genomes project (which corresponds to equal weights). Each vertical line represents a summary of 74 simulated regions (the dot being the median, the line range representing 0.025 to 0.975 quantile). The x-axis shows the three different strategies: using theoretical best possible weights in black (if the estimated effect sizes were to be known), variance matching in green and *Adapt-Mix* in blue. The y-axis shows the MSE-ratio. The MSE-ratio represents the MSE when choosing the weights according to the respective optimisation relative to the MSE when choosing equal weights for all populations (hence a weight of 0.2 for all five populations), i.e. in black h(***w***\*)/*h*(0.2), in green h(***w***^VM^)/*h*(0.2), and in blue h(***w***^AM^)/*h*(0.2). Function *h*(***w***) is the MSE between the estimated and the imputed effect sizes described in Eq. (Eq. (14)). Values on the y-axis smaller than 1 show a smaller MSE in imputation with a specific ***w*** compared to the choice of an unadjusted reference panel with equal weights, while values larger than 1 indicate a higher MSE. Each row represents a subset of different sizes of reference panels, while a msubset of the different Finnish fractions in the GWAS populations are grouped by column. Variants are also grouped according to *F*_*ST*_, with population specific results being on the lower and population unspecific results on upper part of the graph. Figure S3 shows the same graph for all reference panel sizes and GWAS compositions. Table Table 1 provides the same information in a text file. Table B provides the results for all 8 reference panel sizes and fraction of Finnish in the GWAS.

**Table 1.** This table presents the results corresponding to Figure Fig. 1: each entry represents the median MSE (in bold) and the 0.025 - 0.975 quantile in brackets for different Finnish fractions in the GWAS populations are (columns), *F*_*ST*_, different methods (***w***\*, ***w***^VM^ and ***w***^AM^) and reference panel size.

| $F_{ST}$ | Sample size<br>reference panel | Method | Fraction Finnish in GWAS | | |
| --- | --- | --- | --- | --- | --- |
|  |  |  | 20% | 60% | 80% |
| 0-1% | 500 | $w^*$ | **0.996** (0.97-1) | **0.974** (0.925-1) | **0.957** (0.886-0.996) |
| 0-1% | 500 | $w^{VM}$ | **1** (0.996-1.01) | **0.975** (0.929-1.02) | **0.959** (0.89-1.02) |
| 0-1% | 500 | $w^{AM}$ | **1** (0.984-1.02) | **0.985** (0.935-1.03) | **0.966** (0.892-1) |
| 0-1% | 5000 | $w^*$ | **1** (0.974-1) | **0.923** (0.798-0.976) | **0.8** (0.652-0.927) |
| 0-1% | 5000 | $w^{VM}$ | **1** (0.999-1.02) | **0.924** (0.803-0.997) | **0.805** (0.656-0.937) |
| 0-1% | 5000 | $w^{AM}$ | **1** (0.997-1.03) | **0.93** (0.821-0.981) | **0.805** (0.667-0.947) |
| 0-1% | 12500 | $w^*$ | **1** (0.975-1) | **0.891** (0.769-0.966) | **0.742** (0.582-0.884) |
| 0-1% | 12500 | $w^{VM}$ | **1** (0.996-1.02) | **0.892** (0.77-0.977) | **0.747** (0.596-0.895) |
| 0-1% | 12500 | $w^{AM}$ | **1** (0.994-1.03) | **0.9** (0.774-0.969) | **0.751** (0.591-0.913) |
| 1-100% | 500 | $w^*$ | **0.979** (0.781-1) | **0.928** (0.637-1) | **0.886** (0.596-1) |
| 1-100% | 500 | $w^{VM}$ | **1** (0.983-1.02) | **0.937** (0.781-1.1) | **0.892** (0.66-1.12) |
| 1-100% | 500 | $w^{AM}$ | **1** (0.936-1.15) | **0.969** (0.805-1.04) | **0.926** (0.733-1.07) |
| 1-100% | 5000 | $w^*$ | **0.996** (0.906-1) | **0.853** (0.555-0.99) | **0.708** (0.377-0.951) |
| 1-100% | 5000 | $w^{VM}$ | **1** (0.986-1.03) | **0.861** (0.559-1.09) | **0.722** (0.38-1.06) |
| 1-100% | 5000 | $w^{AM}$ | **1** (0.982-1.1) | **0.875** (0.56-1.05) | **0.736** (0.386-1) |
| 1-100% | 12500 | $w^*$ | **0.995** (0.908-1) | **0.814** (0.507-0.975) | **0.648** (0.334-0.908) |
| 1-100% | 12500 | $w^{VM}$ | **1** (0.99-1.03) | **0.83** (0.522-1.03) | **0.658** (0.34-0.951) |
| 1-100% | 12500 | $w^{AM}$ | **1** (0.968-1.08) | **0.845** (0.522-1.01) | **0.658** (0.341-0.944) |

From the extensive simulation results (Fig. 1) it is clear, that the ad-hoc reference panel with equal weights works best (i.e. MSE-ratio close to 1) in two scenarios: for an equally partitioned GWAS (independent of reference panel size or whether the variants are population specific) and when the reference panel is small in size (*n* <= 1000). For all other scenarios, i.e. either *n* > 1000 or the fraction of Finnish in the GWAS is not 0.2, the MSE-ratio is well below 1, therefore indicating a smaller MSE for the optimisation scenario. Note that the *benchmark* MSE-ratio is, by definition, always lower than *Adapt-Mix* and *variance matching* (as it the best theoretically possible MSE).

When comparing *variance matching* and *Adapt-Mix* to the *benchmark* solution, we find that both optimising methods show a similar trend (greater advantage of using specific weights for population specific markers, large reference panel and heterogeneous GWAS). Except for three instances (population non-specific variants when imputed with a reference panel with 5′000 individuals and a GWAS with 40% or 80% Finnish individuals, and specific variants when imputed with a reference panel with 12′500 individuals and a GWAS with 100% Finnish individuals), *variance matching* offers equal or lower MSE than *Adapt-Mix*. When comparing the median MSE-ratio of the optimisation methods to the *benchmark, Adapt-Mix* is performing worst among population specific variants, a reference panel size of 500 and a GWAS with Finnish individuals only. *Variance matching* is performing worst in similar conditions, but when the GWAS consists of no Finnish individuals.

For a reference panel of 500 individuals, population specific variants and a GWAS with 80% Finnish individuals we observe a median MSE-ratio for the theoretically best possible reference panel composition of 0.886, while it is 0.892 for *variance matching* and 0.926 for *Adapt-Mix*. When increasing sample size to 12′500 the MSE-ratio for the *benchmark* solution using becomes 0.648 and 0.658 for *variance matching* and *Adapt-Mix*, respectively.

For variants that are not population specific we see a similar trend with increasing fraction of Finnish individuals and reference panel size, but as expected, less pronounced. For a reference panel of 500 individuals, population unspecific variants and a GWAS with 80% Finnish individuals we observe a median MSE-ratio of 0.957, while it is 0.959 for *variance matching* and 0.966 for *Adapt-Mix*. When increasing sample size to 12′500 it drops to 0.742, 0.747 and 0.751, for ***w***\*, variance matching and *Adapt-Mix*, respectively.

For details to ***w***^VM^, ***w***\* and ***w***^AM^ check Fig. S1.

## 4 Discussion

Summary statistics are used more and more frequently for downstream analyses, but are not always available for all desired variants. These missing summary statistics can, however, be directly imputed from publicly available data using *summary statistics imputation*. The covariance matrices required for this are difficult to estimate from publicly available reference panels due to their size and population structure, requiring their careful adjustment with shrinkage parameters. To address these limitations, we extended the *summary statistics imputation* method as presented in Pasaniuc *et al*. (2014) with an optimal combination of covariance matrices form reference panel subpopulation.

### Choice of reference panel

Formulae for *summary statistics imputation* have two components: GWAS summary statistics and LD matrix estimates which represent the correlation between SNVs. The latter matrix is highly dependent on the reference panel composition: if the ancestry is different between the GWAS and the reference panel, the LD estimation will be biased and yield erroneous summary statistics. An adequate reference panel is therefore critical to the accuracy of *summary statistics imputation*, unlike *genotype imputation* where the Hidden Markov Model makes panel composition much less relevant. Most often, the reference panel for *summary statistics imputation* is often chosen ad-hoc, guessing the underlying GWAS population admixture.

### Variance matching and Adapt-Mix

To tackle this problem, Park *et al*. (2015) proposed *Adapt-Mix* and we propose *variance matching* (Eqs. (12) and (13)). Both methods assume that the GWAS sample is composed of a mixture (or admixture) of populations and that we have a separate reference panel for each population. They then calculate the local LD structure as a linear combination of population-specific estimates, where the weight of each population depends either on the Z-statistics (*Adapt-Mix*) or the allele frequency (*variance matching*). *Variance matching* performed consistently, yet not significantly better than *Adapt-Mix* in 57 out of 60 subgroups that we explored (95% quantile ranges are overlapping in Figure S3). We also found, that *variance matching* offers performance very close to the best possible reference panel composition ***w***\*.

### Variance-bias tradeoff

Although the aim is to approximate the true mixing weights in the GWAS sample, the weights returned will usually deviate from that in an attempt to minimise the MSE. For example, given a GWAS performed in an exclusively Finnish population and using the 1000 Genomes project reference panel, we could either use only the 99 Finnish individuals (weight of 1 for the Finnish population, 0 for others), or select all 503 individuals of the European panel (weight of about 0.2 for the Finnish population). Using only Finnish individuals would more closely match the GWAS allele frequencies and reduce bias, however using the full panel would increase the precision of the estimated correlation matrix, reducing variance (Figure S2). Our approach aims to strike a balance between bias and variance by finding an optimal weight, somewhere between 0.2 and 1 in this example. We find that for smaller reference panels (*n* = 500) the optimal weight tends towards lower values, relying more on information from other populations, whereas for larger panels (*n* = 12′500) the optimal weights tend to be closer to the true underlying population composition in the GWAS (Fig. 2).

**Figure 2.**
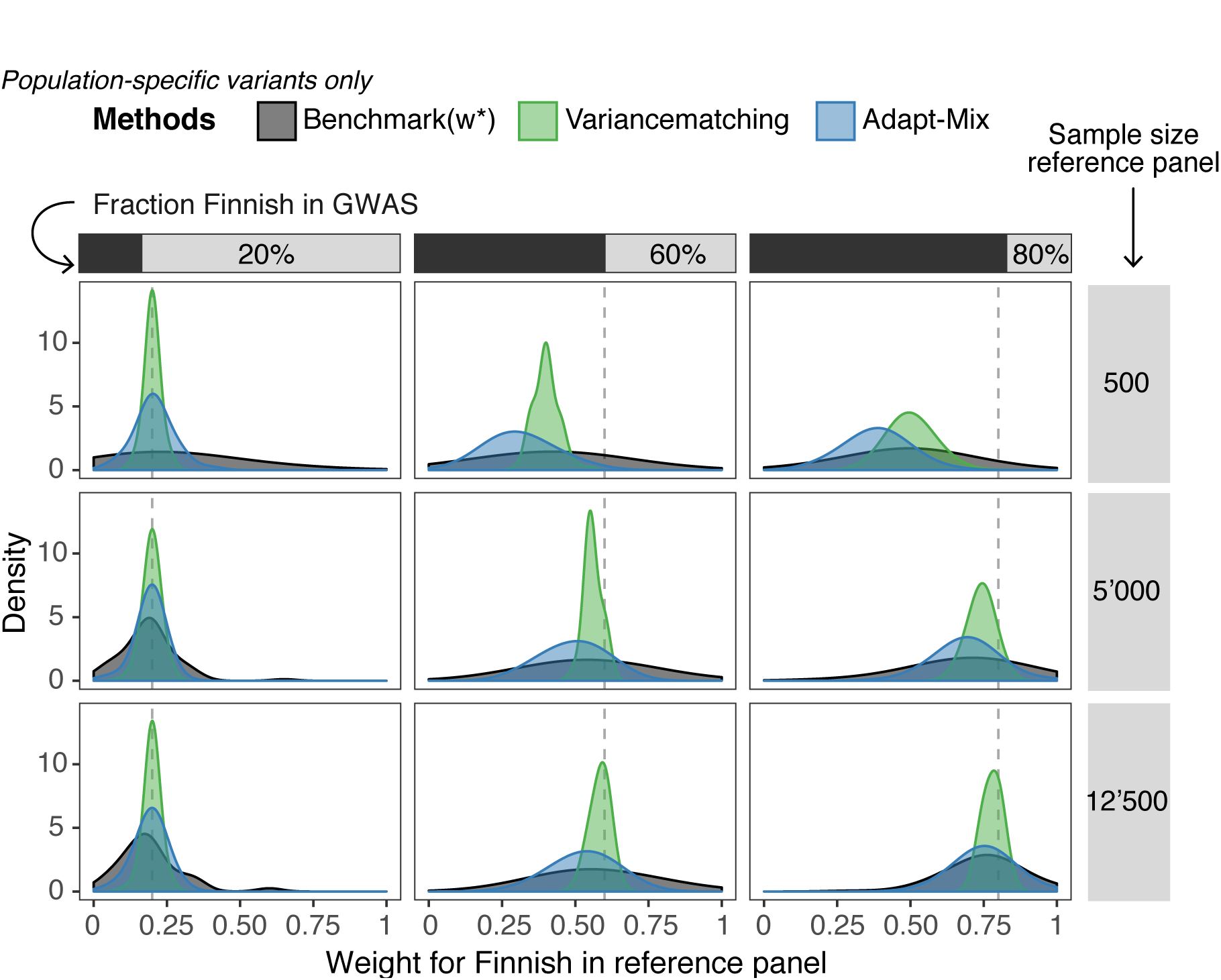
Comparing *w* determined by different methods. This figure compares the weights chosen by all three optimisation methods for population specific variants: ***w***\* as best possible weight (black), ***w***^VM^ by variance matching (green), and ***w***^AM^ by *Adapt-Mix* (blue). ***w***\* represents the benchmark weight: the best possible choice if we were to know the true effect size, but given the same reference panel as for *Adapt-Mix* and variance matching. The x-axis displays the weights for the reference panel chosen by each method, and the y-axis shows the density. The results are split into columns and rows, with the rows for different reference panel sizes and the columns different Finnish fractions in the GWAS populations (also highlighted with the vertical dashed line). Each window contains ***w***\*, ***w***^VM^ and ***w***^AM^ for each of the 74 regions.

### Limitations

*Variance matching assumes* that the population admixture that is reflected in the variance of tag SNVs (diagonal in matrix ***C***) is the same as the covariance between tag SNVs (off-diagonal of *C*) as well as between tag SNVs and SNVs to impute (matrix c). Furthermore, our analytical solution to Eq. (11) involves approximations of the variance and the bias (Eq. (S7) and (S8)).

In general, finding a reference panel whose ancestry composition matches that of the GWAS is difficult because the mixture/admixture of populations is usually unknown. With *variance matching* we are addressing this by composing a matching LD matrix. However, there are other challenges too: publicly available reference panels have a limited number of populations with a limited number of individuals. To this end, we could not validate our approach in real data as diverse reference panels with sample sizes > 500 per population are not publicly available at this time.

Due to lack of large, sequenced reference panels we used an upsampling technique called HAP-GEN2 (Su *et al*., 2011), which limits the lower bound of the global allele frequency to 1/(2 · 503). Finally, the outcome used for the simulated GWAS is based on one causal variant with an explained variance of 0.02, therefore it might not be fully representative for a polygenic phenotype with more than one causal variant.

Finally, our method is not applicable to GWAS studies that decided not to share allele frequency information.

## 5 Conclusion

With variance matching we present an extension to the published summary statistics imputation method (Pasaniuc *et al*., 2014) by allowing the LD structure to be adaptively estimated according to population admixture. To evaluate this extension, we performed GWAS on upsampled 1000 Genomes project data in combination with a simulated phenotypes. Due to the bias-variance trade-off, accounting for differences in population admixture between GWAS and reference panel yields better results with increasing panel size.

## Acknowledgements

Ninon Mounier, Anthony Sonrel and Jonathan Sulc gave valuable comments on an earlier draft of the manuscript.

## Funding

This work was supported by the Leenards Foundation (http://www.leenaards.ch), the Swiss Institute of Bioinformatics (http://www.isb-sib.ch/), and the Swiss National Science Foundation [31003A-143914, 31003A-169929].

## References

1000 Genomes Project Consortium (2010). A map of human genome variation from population-scale sequencing. Nature, 467(7319), 1061–1073.

Bulik-Sullivan, B., Finucane, H., Anttila, V., Gusev, A., Day, F., Loh, P., ReproGen Consortium, Psychiatric Genomics Consortium, Genetic Consortium for Anorexia Nervosa of the Wellcome Trust Case Control Consortium 3, Duncan, L., and Perry, J. (2015). An atlas of genetic correlations across human diseases and traits. Nature Genetics, 47(11), 1236–1241.

Burgess, S., Butterworth, A., and Thompson, S. G. (2013). Mendelian randomization analysis with multiple genetic variants using summarized data. Genetic Epidemiology, 37(7), 658–665.

Eaton, M. L. (1983). Multivariate Statistics: A Vector Space Approach. John Wiley & Sons Inc.

Lee, D., Bigdeli, T. B., Riley, B. P., Fanous, A. H., and Bacanu, S. A. (2013). DIST: Direct imputation of summary statistics for unmeasured SNPs. Bioinformatics, 29(22).

Lee, D., Williamson, V. S., Bigdeli, T. B., Riley, B. P., Fanous, a. H., Vladimirov, V. I., and Bacanu, S.-A. (2014). JEPEG: a summary statistics based tool for gene-level joint testing of functional variants. Bioinformatics, 31(8).

Lee, D., Bigdeli, T. B., Williamson, V. S., Vladimirov, V. I., Riley, P., Fanous, A. H., and Bacanu, S.-a. (2015a). Genome analysis distmix: Direct imputation of summary statistics for unmeasured snps from mixed ethnicity cohorts. Bioinformatics.

Lee, D., Williamson, V. S., Bigdeli, T. B., Riley, B. P., Webb, B. T., Fanous, A. H., Kendler, K. S., Vladimirov, V. I., and Bacanu, S.-A. (2015b). JEPEGMIX: gene-level joint analysis of functional SNPs in cosmopolitan cohorts: Table 1. Bioinformatics, 32.

Lim, E. T., Würtz, P., Havulinna, A. S., Palta, P., Tukiainen, T., Rehnström, K., Esko, T. o., Mägi, R., Inouye, M., Lappalainen, T., Chan, Y., Salem, R. M., Lek, M., Flannick, J., Sim, X., Manning, A., Ladenvall, C., Bumpstead, S., Hämäläinen, E., Aalto, K., Maksimow, M., Salmi, M., Blankenberg, S., Ardissino, D., Shah, S., Horne, B., McPherson, R., Hovingh, G. K., Reilly, M. P., Watkins, H., Goel, A., Farrall, M., Girelli, D., Reiner, A. P., Stitziel, N. O., Kathiresan, S., Gabriel, S., Barrett, J. C., Lehtimäki, T., Laakso, M., Groop, L., Kaprio, J., Perola, M., McCarthy, M. I., Boehnke, M., Altshuler, D. M., Lindgren, C. M., Hirschhorn, J. N., Metspalu, A., Freimer, N. B., Zeller, T., Jalkanen, S., Koskinen, S., Raitakari, O., Durbin, R., MacArthur, D. G., Salomaa, V., Ripatti, S., Daly, M. J., and Palotie, A. (2014). Distribution and Medical Impact of Loss-of-Function Variants in the Finnish Founder Population. PLoS Genetics, 10(7).

Marchini, J. and Howie, B. (2010). Genotype imputation for genome-wide association studies. Nature Reviews Genetics, 11(7), 499–511.

McEvoy, B. P., Montgomery, G. W., McRae, A. F., Ripatti, S., Perola, M., Spector, T. D., Cherkas, L., Ahmadi, K. R., Boomsma, D., Willemsen, G., Hottenga, J. J., Pedersen, N. L., Magnusson, P. K. E., Kyvik, K. O., Christensen, K., Kaprio, J., Heikkilä, K., Palotie, A., Widen, E., Muilu, J., Syvänen, A. C., Liljedahl, U., Hardiman, O., Cronin, S., Peltonen, L., Martin, N. G., and Visscher, P. M. (2009). Geographical structure and differential natural selection among North European populations. Genome Research, 19(5), 804–814.

Park, D. S., Brown, B., Eng, C., Huntsman, S., Hu, D., Torgerson, D. G., Burchard, E. G., and Zaitlen, N. (2015). Adapt-Mix: learning local genetic correlation structure improves summary statistics-based analyses. Bioinformatics, 31(12).

Pasaniuc, B., Zaitlen, N., Shi, H., Bhatia, G., Gusev, A., Pickrell, J., Hirschhorn, J., Strachan, D. P., Patterson, N., and Price, A. L. (2014). Fast and accurate imputation of summary statistics enhances evidence of functional enrichment. Bioinformatics, 30(20).

Su, Z., Marchini, J., and Donnelly, P. (2011). HAPGEN2: simulation of multiple disease SNPs. Bioinformatics.

Wen, X. and Stephens, M. (2010). Using linear predictors to impute allele frequencies from summary or pooled genotype data. Annals of Applied Statistics, 4(3), 1158–1182.

Xu, Z., Duan, Q., Yan, S., Chen, W., Li, M., Lange, E., and Yun, L. (2015). Genome Analysis DISSCO: Direct Imputation of Summary Statistics allowing COvariates. Bioinformatics.

Yang, J., Ferreira, T., Morris, A. P., Medland, S. E., Madden, P. a. F., Heath, A. C., Martin, N. G., Montgomery, G. W., Weedon, M. N., Loos, R. J., Frayling, T. M., McCarthy, M. I., Hirschhorn, J. N., Goddard, M. E., and Visscher, P. M. (2012). Conditional and joint multiple-SNP analysis of GWAS summary statistics identifies additional variants influencing complex traits. Nature Genetics, 44(4), 369–375.

